# Tissue mechanics controls T-cell activation and metabolism

**DOI:** 10.1101/581322

**Authors:** Kevin P. Meng, Fatemeh S. Majedi, Timothy J. Thauland, Manish J. Butte

**Affiliations:** Department of Microbiology and Immunology, Stanford University, Stanford, California 94305, USA.; Department of Bioengineering, University of California, Los Angeles, Los Angeles, CA, 90095, USA.; Division of Immunology, Allergy, and Rheumatology, Department of Pediatrics, University of California Los Angeles, Los Angeles, CA, 90095, USA.; Department of Microbiology, Immunology, and Molecular Genetics, University of California Los Angeles, Los Angeles, CA, 90095, USA.

## Abstract

Upon immunogenic challenge, lymph nodes become mechanically stiff as immune cells proliferate within their encapsulated environments, and with resolution, they reestablish a soft, baseline state. We found that these mechanical changes in the microenvironment promote and then restrict T-cell activation and metabolic reprogramming. Sensing of tissue mechanics by T cells requires the mechanosensor YAP. Unlike in other cells where YAP promotes proliferation, YAP in T cells suppresses proliferation in a stiffness-dependent manner by directly restricting the translocation of NFAT into the nucleus. YAP regulates T-cell responses against viral infections and in autoimmune diabetes. Our work reveals a new paradigm whereby tissue mechanics fine-tunes adaptive immune responses in health and disease.

**One Sentence Summary:** Tissue mechanics regulates T cells.

## Main Text

T cells encounter forces at the tissue scale as they traverse through a variety of microenvironmental mechanical contexts, including sites of infection, inflammation, and cancer. T cells are also sensitive to forces at the molecular scale, as triggering of the T cell receptor itself requires forces^1–3^ that arise through a series of pushing and pulling movements^3,4^ that correspond to spreading of the T cell upon an APC^5,6^. Activation of the TCR is potentiated when the cognate peptide-MHC (or monoclonal antibody to CD3) is anchored to stiff antigen presenting cells, stiff artificial pillars, or stiff polymer matrices^7–10^. However, mechanosensation of the tissue microenvironment differs from the mechanical forces exerted on the TCR, as tissue-scale forces can be sensed through a variety of receptors or directly by the cytoskeleton^11^. The transcriptional coactivator Yes-associated protein (YAP) is a well-known sensor of tissue mechanics that is dominantly regulated by the stiffness of extracellular matrix^12^. Understanding microenvironmental mechanosensing has offered revolutionary leaps in the fields of cellular development, organ size regulation, stem cell differentiation, cancer metastasis, cellular senescence, and programmed cell death^13–16^. Recent works have established a role for YAP in immunity, but there have been no links yet between YAP and mechanics in immune cells^17–20^, and environmental mechanosensing has never been demonstrated for T cells.

To understand the impact of forces at the tissue scale, we studied lymph nodes (LNs) during infection. Activated LN become stiff and swollen during an immune response, but shrink and return to a baseline of mechanical softness upon resolution. Thus, the rise and fall of the adaptive immune response appears to synchronize with changes in LN stiffness. To shed light on T-cell pathophysiology in disease-associated tissues, we sought to determine how mechanical signals translate to regulation of the T-cell effector program.

## RESULTS

### Tissue mechanics regulate T cell proliferation and activation

Infections have been noted for centuries to cause inflammation and induce rigidity of regional lymph nodes. To quantitate the changes in lymph node stiffness over the course of an infection, we measured lymph nodes of mice infected with lymphocytic choriomeningitis virus (LCMV) Armstrong. We observed a ~10-fold increase of stiffness (elastic modulus) at day 3 and 7, after which the elastic modulus returned back towards baseline (**Fig. 1A**). The size of the LN matched these mechanical changes (**Fig. S1A**). To assess the effects of stiffness on T-cell activation, we generated biomimetic alginate-RGD scaffolds of 4 kPa (soft) and 40 kPa (stiff) elastic moduli that recapitulated the lymph node mechanics observed *in vivo* (**Fig. 1B**, **Fig. S1B-E**). SMARTA TCR transgenic CD4+ T cells were co-cultured with syngeneic splenocytes atop these scaffolds and stimulated with their cognate peptide GP 61-80 for 2-4 days. T cells showed increased proliferation when cultured on the stiffer substrate as compared to the softer substrate, reducing the dose of antigen required for 50% of maximal proliferation (ED50) by ~2-fold (**Fig. 1C**). Activating T cells on stiffer surfaces also led to enhanced activation markers, including CD25 and CD44 (**Fig. 1D**). Thus, mechanically stiff microenvironments, like that of activated lymph nodes, augment T-cell activation.

**Fig. 1.**
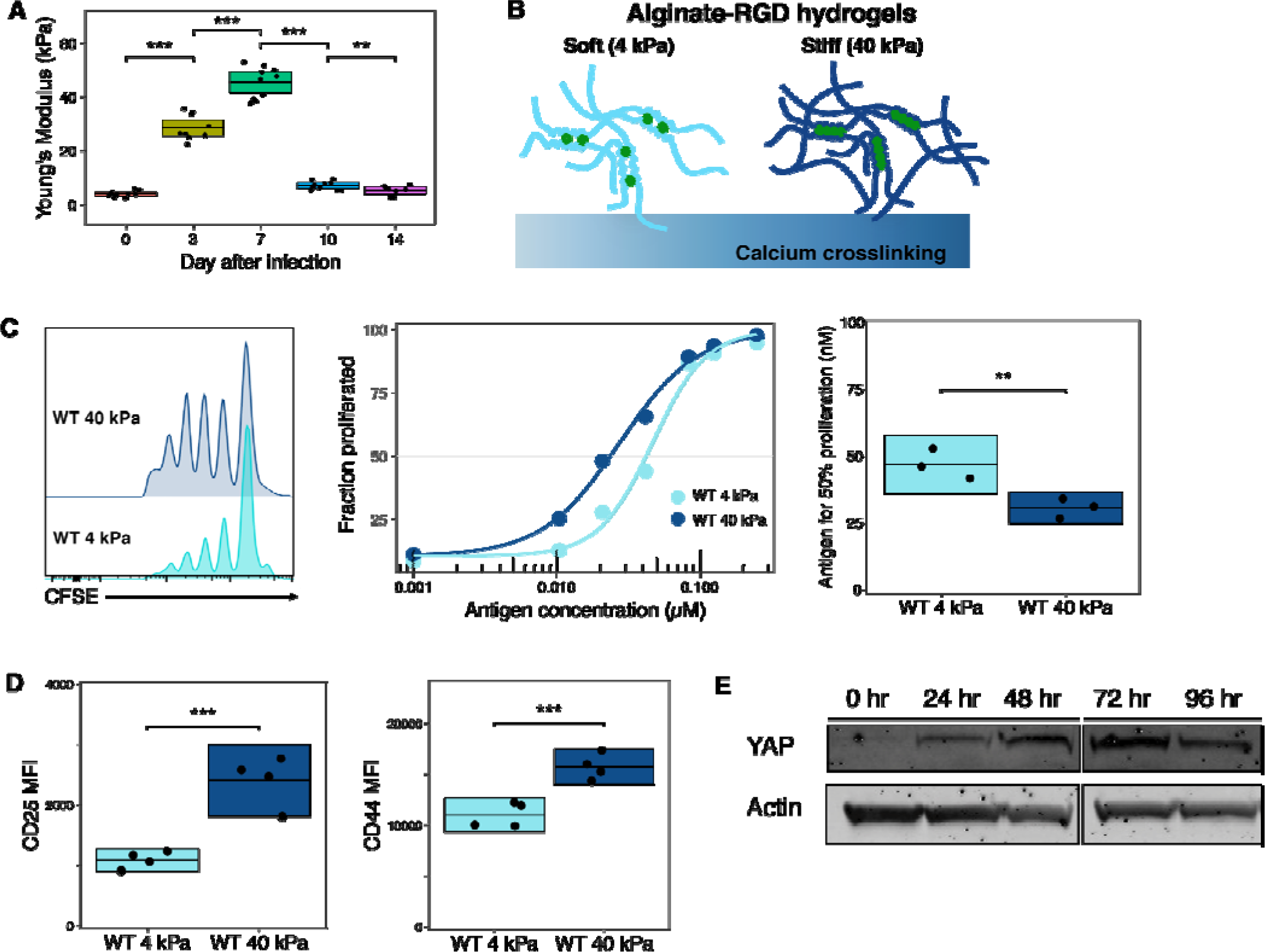
Tissue mechanics regulate T cell proliferation and activation. (A) Inguinal and brachial lymph nodes from LCMV-infected mice measured for mechanical stiffness. Each point represents one lymph node. 2-3 lymph nodes were measured from each mouse × 3 mice at each timepoint. Data representative of 3 independent experiments. Box shows mean and 95% CI. (B) Schematic of 4 kPa and 40 kPa 2D scaffolds generated by alginate-RGD crosslinked with calcium. (C) Proliferation (left) of CFSE-labeled SMARTA T cells cultured onto soft (light blue) or stiff (dark blue) scaffolds with splenocytes and GP 61-80 peptide. ED50 calculated from three such independent dose-response experiments shown (right). Box shows mean and 95% CI. (D) T-cell activation markers after 72 h stimulation on soft (light blue) or stiff (dark blue) scaffolds with APCs plus 83 nM GP 61-80 peptide. Data from four independent experiments are shown. Box shows mean and 95% CI. (E) Immunoblot for YAP in mouse CD4+ T cells stimulated at different timepoints. Data representative of 3 independent experiments.

The transcription coactivator YAP has been shown to play important roles in mechanosensing, growth, and differentiation of assorted cell types. However, its role in T cells remains largely unknown. To test whether YAP is expressed in CD4+ T cells, we measured protein expression in T cells from WT mice *ex vivo* and after stimulation. We discovered that YAP is dramatically induced upon T-cell activation (**Fig. 1E**). We also observed increased nuclear localization of YAP in T cells experiencing stiff surfaces (**Fig. S1F**). These results show that T cells sense their mechanical microenvironment, that YAP is induced upon T-cell activation, and that YAP traverses into the nucleus in a stiffness-dependent manner.

### YAP restricts TCR-induced CD4+ T cell activation and proliferation

We sought to determine if YAP played a role in the sensing of the mechanical microenvironment in T cells, a question that will be addressed below, but first we developed an approach to study this question. We created a conditional knockout of YAP (YAPcKO) by crossing YAP flox mice to CD4-cre mice. YAP protein was absent in both CD4+ and CD8+ effector T cells (**Fig. S2A**). In other cell types, the Mst1/2 kinases are important regulators of YAP. Mst1/2 conditional knockout mice displayed defects in thymic egress of T cells ^21^, but we did not observe any developmental defects in thymic or splenic T cells in YAPcKO mice (**Fig. S2B-S2E**). Furthermore, we found no difference in percentages of natural Foxp3+ Treg cells (**Fig. S2F**), or in skewed development of iTreg or Th17 cells (**Fig. S2F-S2G**) in comparing YAPcKO and control littermates.

Given that YAP protein is highly upregulated in effector T cells, we hypothesized that YAP was important for effector T-cell responses. Surprisingly, activating YAPcKO CD4+ T cells led to an increase in the proliferation compared to that of WT CD4+ T cells, with a decrease in ED50 by 1.5-fold (**Fig. 2A**). Activated YAPcKO T cells also secreted more IL-2 (**Fig. 2B**) and showed a marked increase in activation markers, including CD69, CD25, and CD44 (**Fig. 2C**). To confirm that YAP negatively regulates T-cell proliferation and activation, we performed a “rescue” experiment. We transduced the full-length cDNA for YAP (FL YAP) or a control vector into TCR transgenic OT-II YAPcKO T cells, then re-stimulated the transduced T cells with APCs plus a range of cognate ovalbumin peptide. YAP expression in YAPcKO T cells led to decreased proliferation, IL-2 production, and CD25 expression compared to vector control (**Fig. 2D-2F**). Together, these data establish YAP as a suppressor of T-cell activation and effector responses.

**Fig. 2.**
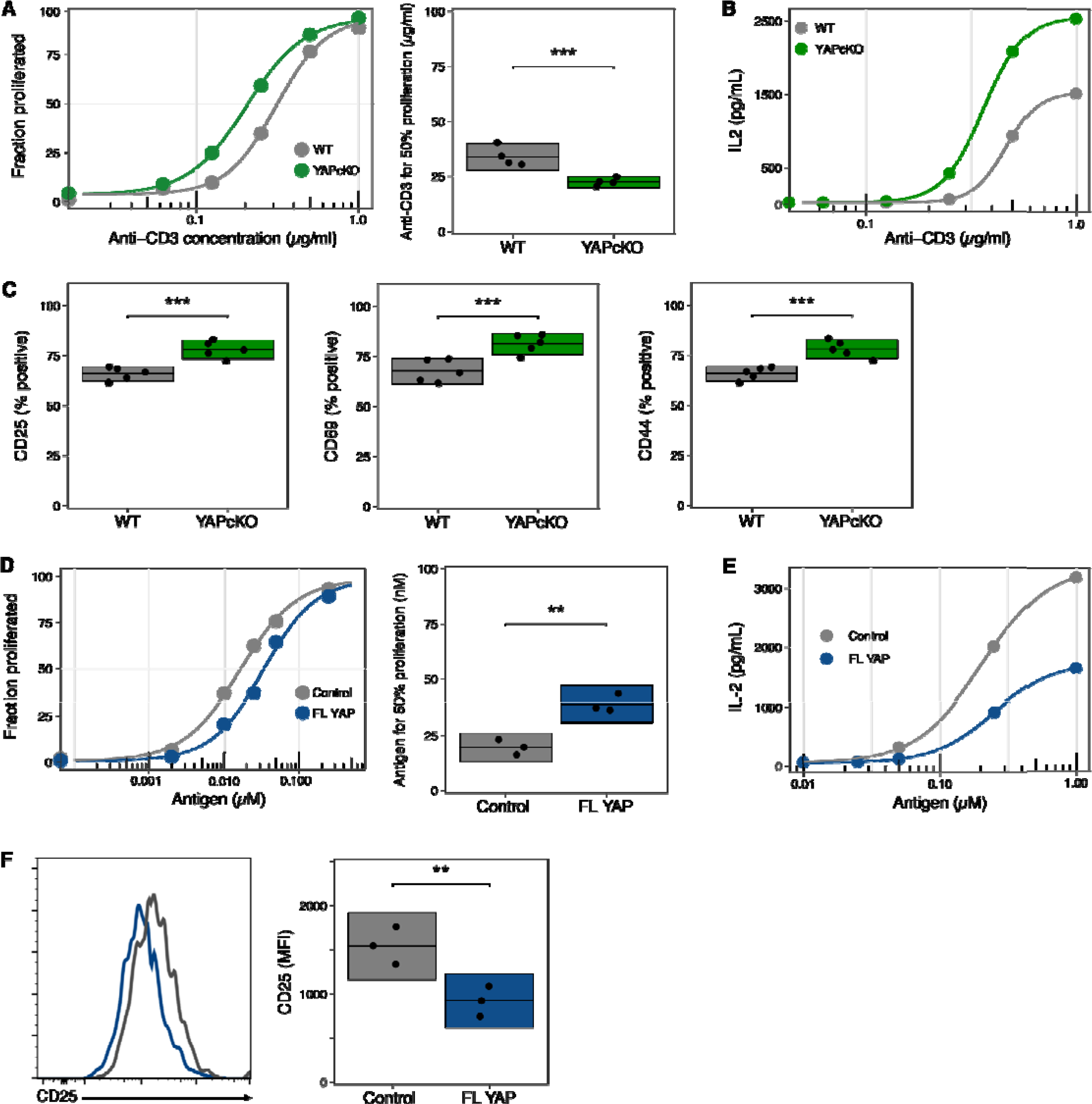
YAP restrains T cell proliferation and effector function. (A) Proliferation (left) of CFSE-labeled WT (gray) or YAPcKO (green) T cells after anti-CD3 and anti-CD28 stimulation. ED50 calculated from four such independent dose-response experiments shown (right). Box shows mean and 95% CI. (B) IL-2 secretion of WT or YAPcKO T cells at 72 h after activation with anti-CD3 and anti-CD28 stimulation. Data representative of three independent experiments. (C) Activation markers of WT or YAPcKO T cells stimulated with 1 µg/ml anti-CD3 for 24 hours. Data from five independent experiments are shown. Box shows mean and 95% CI. (D) Proliferation of CellTrace Violet-labeled OT-II YAPcKO T cells transduced with control vector (gray) or FL YAP (blue) after restimulation with OVA 323-339 peptide. ED50 calculated from three such independent dose-response experiments shown (right). Box shows mean and 95% CI. (E) IL-2 secretion of OT-II YAPcKO T cells transduced with control vector (gray) or FL YAP (blue) at 48 hours post restimulation with OVA peptide. Data representative of two independent experiments. (F) CD25 expression of restimulated T cells from experiment in (E). Data from three independent experiments are shown. Box shows mean and 95% CI.

### YAP inhibits antiviral T-cell effector response

YAP has been implicated in the innate immune response to viral infection ^19^. Thus, we wondered whether YAP could influence adaptive immunity in viral infections. Using an adoptive transfer mouse model, we co-injected equal numbers of congenic, naïve WT and YAPcKO SMARTA T cells, and then infected the naïve recipient mice with a non-lethal dose of LCMV Armstrong (**Fig. 3A**). We observed a significant proliferative advantage of YAPcKO SMARTA T cells versus control SMARTA T cells near the height of the response (**Fig. 3B**), along with heightened activation markers and cytokine production (**Fig. 3C** and **Fig. S3A-B**). These results show a cell-intrinsic effect of YAP on suppressing T-cell effector responses to a viral infection.

**Fig. 3.**
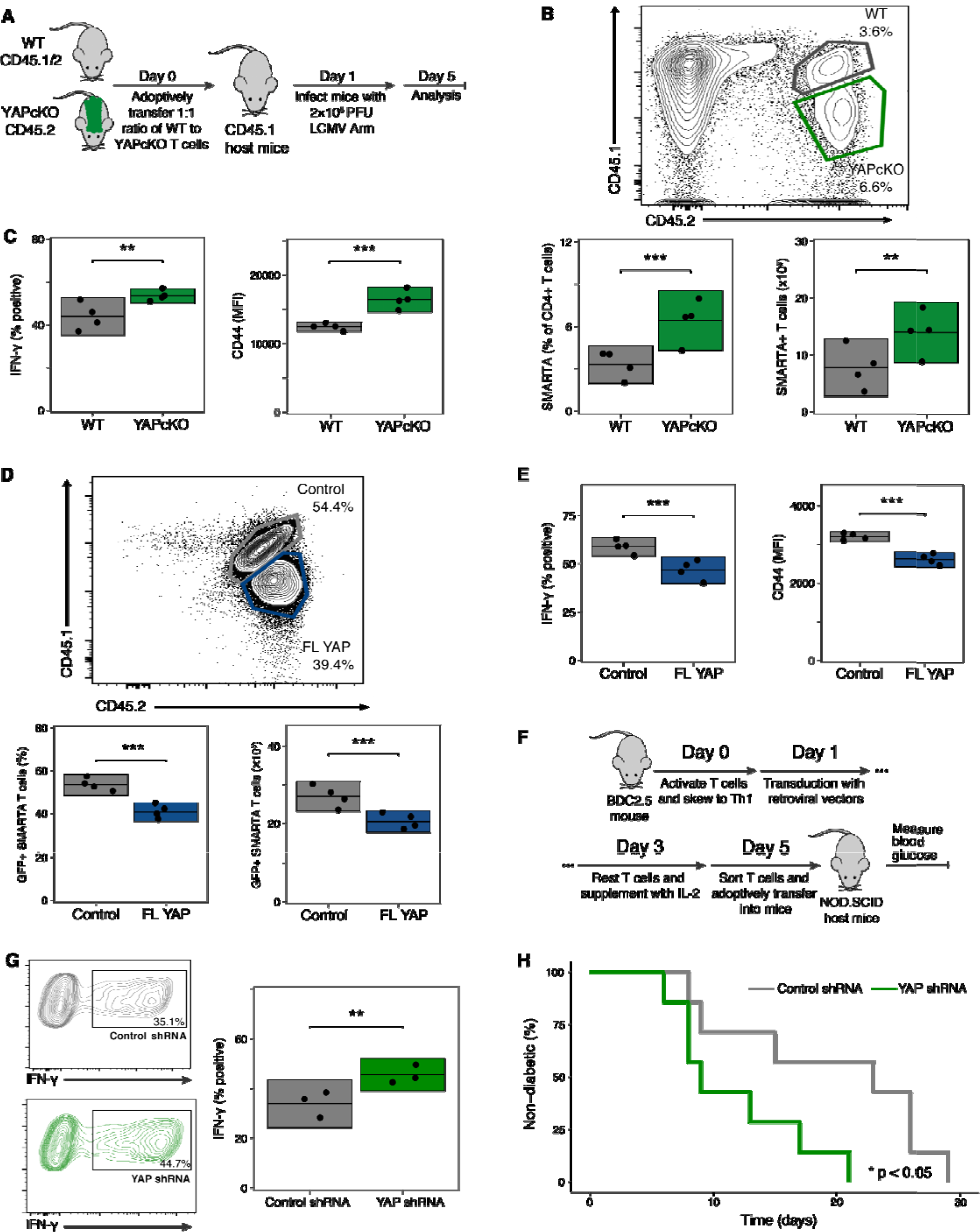
YAP suppresses antiviral response and restricts autoimmunity. (A) Schematic of LCMV adoptive transfer model. (B) The frequency of adoptively transferred SMARTA WT (gray) and YAPcKO T cells (green) in mice on day 5 after infection with LCMV. Data are representative of four independent experiments (n=4 in each group in each experiment). Each dot is one mouse within an experiment, box shows mean and 95% CI. (C) Quantification of CD44 MFI and IFN-□ percentage in SMARTA T cells from experiment in (B). (D) The frequency of adoptively transferred SMARTA YAPcKO T cells transduced with control vector (gray) or FL YAP (blue). Data are representative of three independent experiments (n=4 in each group in each experiment). Each dot is one mouse within an experiment, box shows mean and 95% CI. (E) Quantification of CD44 MFI and IFN-□ percentage in SMARTA T cells from experiment in (D). (F) Schematic of BDC2.5 type I diabetes adoptive transfer model (G) Percentage of restimulated BDC2.5 T cells producing IFN-□ after knockdown with control shRNA (gray) or YAP shRNA (green) in vitro. Data from three independent experiments are shown (right). Box shows mean and 95% CI. (H) Survival curve of NOD.SCID mice receiving T cells expressing either control or YAP shRNA (n = 7 mice per group). Representative of three independent diabetes experiments.

To test whether YAP expression would “rescue” the observed phenotype, we expressed FL YAP or a control vector in YAPcKO SMARTA T cells and co-adoptively transferred them into CD45.1 mice infected with LCMV. We found that over-expression of YAP led to impaired T-cell expansion as compared to those receiving a control vector (**Fig. 3D**), consistent with our *in vitro* data, along with decreased IFN-□ production and CD44 expression (**Fig. 3E**). These results demonstrate that expression of FL YAP rescues the phenotype in YAP-deficient T cells. Thus, YAP inhibits the T-cell response to a viral infection.

### YAP inhibits metabolic reprogramming of activated T cells

In other cell types, YAP plays a role downstream of metabolic sensors to allow proliferation only when there are ample metabolic resources ^22^. YAP activity can also regulate the metabolic state of a cell to support a proliferative program ^23,24^. The metabolic state of T cells is tightly regulated and central to their role in the immune response. To determine whether a metabolic link exists between YAP and T cell effector functions, we used extracellular flux analysis to measure the levels of glycolysis and oxidative phosphorylation (OXPHOS) in WT and YAPcKO T cells. We observed an increase in glycolytic capacity in activated YAPcKO T cells as compared to littermate controls (**Fig. 4A**). There were no differences in basal glucose uptake or basal glycolysis (**Fig. S4B**), which was confirmed using the fluorescent glucose analog 2-NBDG (**Fig. S4A**). Meanwhile, YAPcKO T cells exhibited a greater basal level of oxygen consumption rate (OCR) and maximal respiration capacity compared to WT cells (**Fig. S4C**). Absence of YAP in T cells also led to elevated spare respiratory capacity (SRC), an indicator of the mitochondrial ability to respond to stress ^25^ (**Fig. 4B**). Effector WT and YAPcKO T cells were notably different in regards to basal ECAR/OCR ratio and in their maximal glycolytic and respiratory capacity (**Fig. S4E**). On the other hand, we saw the basal ECAR and OCR of naïve cells were nearly identical, consistent with YAP not being expressed in naïve T cells (**Fig. S4D and S4E**). We found an increase in mitochondrial staining in YAPcKO effector T cells compared to WT T cells (**Fig. 4C**), which could explain the increased respiratory capacity. We also found a higher mitochondrial membrane potential in YAPcKO effector T cells compared to that of WT T cells (**Fig. 4D**), which has been linked to increased ATP production and T-cell effector function ^26^. Thus, YAP plays a role upstream of the metabolic program of effector T cells.

**Fig. 4.**
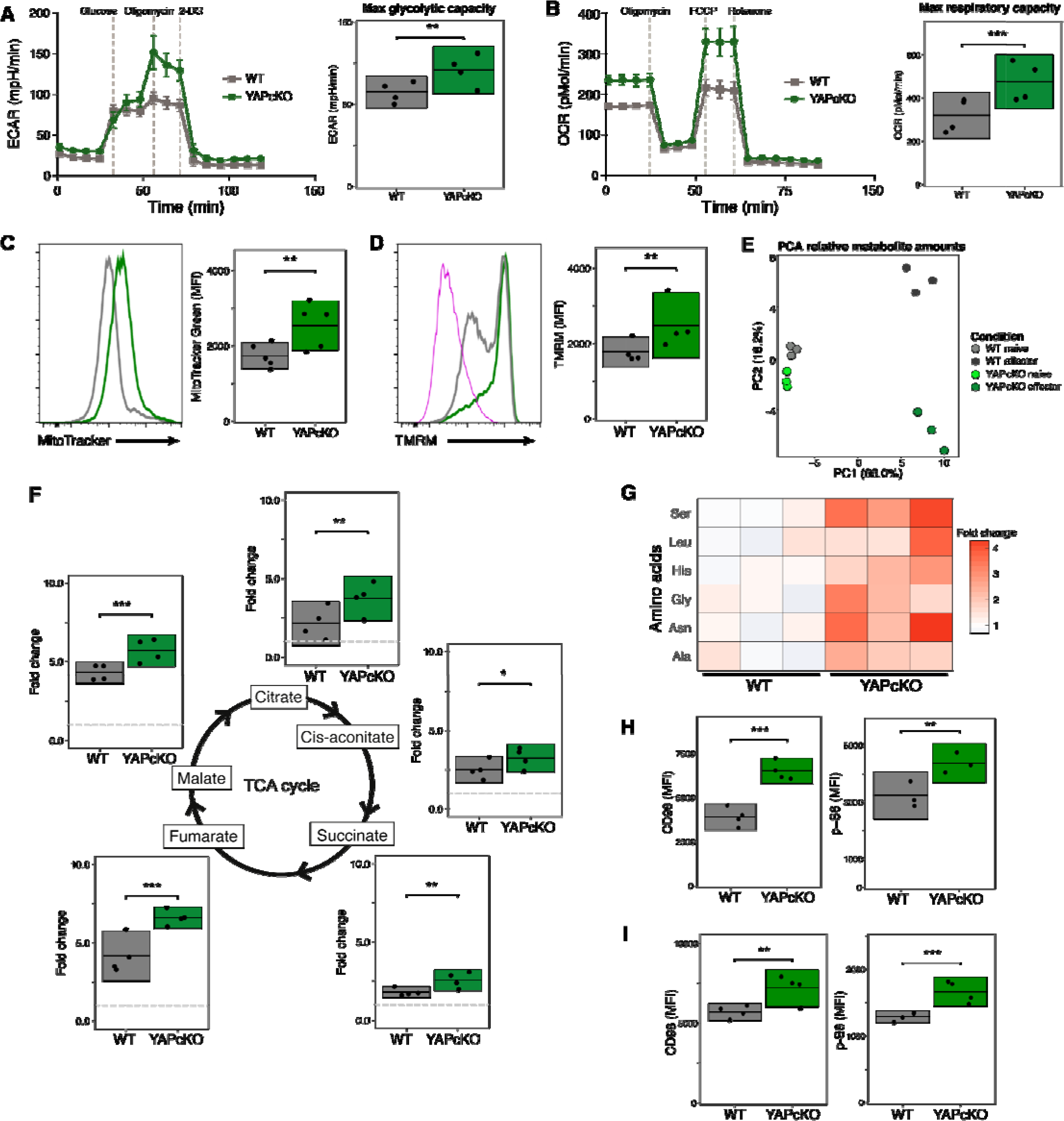
YAP coordinates metabolic reprogramming and amino acid metabolism. (A) Extracellular acidification rate (ECAR) of WT (gray) and YAPcKO (green) effector T cells on day 3 after activation under basal conditions and in response to glucose, oligomycin, and 2-DG. Four such independent experiments are shown, box shows mean and 95% CI (right) of maximal glycolytic capacity. (B) Oxygen consumption rate (OCR) of WT (gray) and YAPcKO (green) effector T cells on day 3 after activation under basal conditions and in response to oligomycin, FCCP, and rotenone/antimycin A. Four such independent experiments are shown, box shows mean and 95% CI (right) of maximal respiratory capacity. (C) WT or YAPcKO effector T cells were analyzed for mitochondria size with MitoTracker Green. Data from five independent experiments are shown. Box shows mean and 95% CI. (D) WT or YAPcKO effector T cells analyzed for mitochondria membrane potential measured using TMRM. FCCP (magenta) was used as a control to ensure mitochondria were depolarized. Data from four independent experiments are shown. Box shows mean and 95% CI. (E) Metabolomics of naive and effector WT/YAPcKO T cells, principle component analysis (PCA) shows relationships among WT and YAPcKO T cells. Data shown are from a collection of three independent experiments. (F) Fold change of intracellular TCA metabolites measured by MS between effector WT and YAPcKO T cells normalized to naive WT or YAPcKO T cell levels (gray dotted line), respectively. Data from four independent experiments are shown. Box shows mean and 95% CI. (G) Heatmap of intracellular amino acid abundance. Intracellular amino acid abundance in WT or YAPcKO effector T cells were first normalized to their respective naïve counterparts. Data are shown as fold change of intracellular amino acid abundance in WT or YAPcKO effector T cells normalized to the average abundance of each amino acid in WT T cells from three independent experiments. (H) CD98 and p-S6 expression in WT (gray) and YAPcKO (green) effector T cells, showing four independent experiments (three for p-S6). Box shows mean and 95% CI. (I) CD98 and p-S6 expression in adoptively transferred WT and YAPcKO SMARTA T cells in mice five days after infection with LCMV. WT (gray) and YAPcKO (green) effector T cells. Data are representative of three independent experiments (n=4 in each group in each experiment). Each dot is one mouse within an experiment, box shows mean and 95% CI.

To gain further insight into the metabolic regulation, we measured polar metabolite levels in cultured T cells. WT and YAPcKO effector T cells showed large differences, especially as compared to naïve T cells (**Fig. 4E**). Metabolites that separate WT and YAPcKO effector T cells included the ADP/ATP and AMP/ATP ratios, amino acids, citric acid cycle intermediates, and nucleotides (**Fig. S4G**). We also observed significant products of metabolic activity, including elevated intracellular lactate and ATP (**Fig. S4F**). The citric acid (TCA) cycle fuels OXPHOS, and so we examined TCA cycle intermediates and found a substantial increase in YAPcKO T cells as compared with controls (**Fig. 4F**). We wondered if YAPcKO effector T cells had more TCA cycle intermediates because of an increased rate of TCA cycling. To address this question, we used a metabolic tracing approach. We incubated effector T cells with ^13^C-glucose or ^13^C-glutamine and measured differences in mass isotopomer distributions between activated WT and YAPcKO T cells. We did not detect a significant difference in the isotopomeric labeling of citrate for ^13^C-glucose-treated or ^13^C-glutamine-treated samples, indicating that YAPcKO T cells had similar rates of TCA cycling to WT T cells (**Fig. S4H**). Indeed, the fractional contribution of labeled glucose to isotopologues of TCA metabolites was mostly similar between WT and YAPcKO T cells (**Fig. S4I**). These labeling results suggest that YAPcKO T cells have more mitochondrial biogenesis rather than enhanced mitochondrial function. Altogether, these data indicate that YAP inhibits pathways of mitochondrial metabolism.

### YAP coordinates mTOR activity and amino acid metabolism

Our metabolomics data revealed that YAPcKO effector T cells had higher intracellular amounts of several amino acids, including serine, glycine, asparagine, and leucine, all of which have been implicated in T-cell effector function and proliferation (**Fig. 4G** and **Fig. S5A**) ^27,28^. In line with the intracellular data, the abundance of these amino acids in the media was markedly reduced in the supernatant of cultured YAPcKO T cells as compared with WT controls (**Fig. S5B**). CD98 (SLC3A2) forms a complex with LAT1 (SLC7A5) for the import of several amino acids, including leucine. Staining for cell-surface CD98 revealed that it was more highly expressed in YAPcKO effector T cells than WT controls (**Fig. 4H**). These data show that during T-cell activation, YAPcKO T cells imported more amino acids from the extracellular environment. Additionally, amino acid metabolism has been closely linked to mTOR signaling ^29^. To test the involvement of mTOR in YAP-mediated effects on T cells, we examined downstream targets of mTOR in effector T cells. We found phosphorylation of S6 (p-S6) was greater in YAPcKO effector T cells compared to that of WT cells (**Fig. 4H**). To test whether elevated amino acid transport and mTOR activity also held true in an *in vivo* setting, we measured the expression of CD98 and p-S6 in adoptively co-transferred SMARTA T cells (WT and YAPcKO) after 5 days of infection with LCMV. YAPcKO SMARTA effector T cells exhibited greater CD98 and p-S6 expression than controls (**Fig. 4I**). Overall, our data show YAP dampens the T-cell response by downregulating mTOR signaling, and amino acid metabolism and transport.

### YAP inhibits NFAT activity and translocation into the nucleus

Regulation of many metabolic changes in effector T cells have largely been attributed to transcription driven by the Nuclear Factor of Activated T cells (NFAT) ^30,31^. Furthermore, store-operated calcium entry and calcineurin regulate mitochondrial activity in an NFAT-dependent manner ^32^. Upon triggering of the T-cell receptor, NFAT1 and NFAT2 travel into the nucleus and drive transcription, one of the targets being IL-2. Given the metabolic changes and the increased IL-2 expression of YAPcKO T cells, we studied the activation of NFAT1 and NFAT2. YAP-deficient effector T cells exhibited greater NFAT1 nuclear localization as compared with WT controls, but not NFAT2 (**Fig. 5A** and **S6A**). We sought next to understand how YAP controls NFAT localization.

**Fig. 5.**
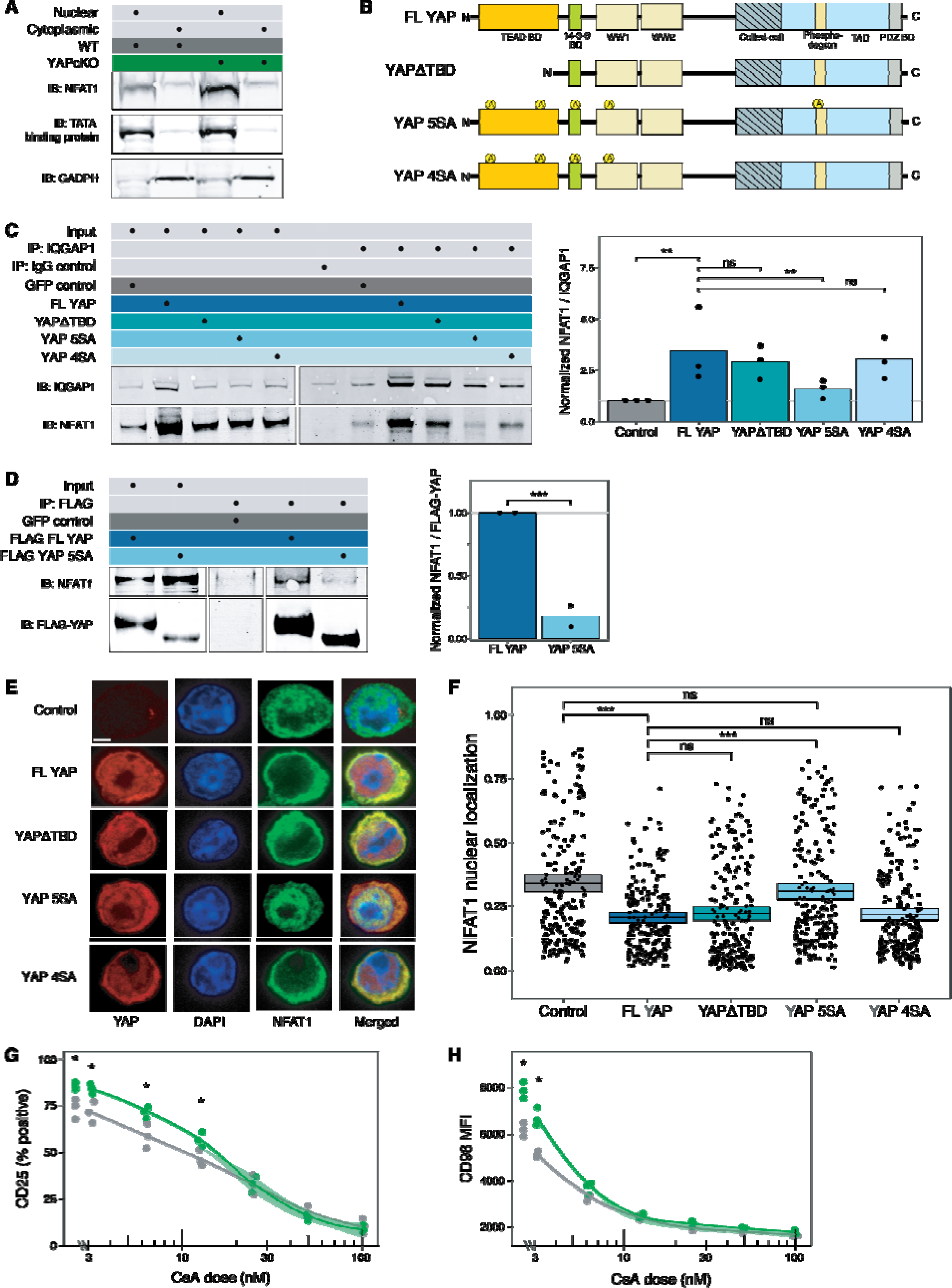
YAP sequesters NFAT1 in the cytoplasm. (A) Immunoblot analysis for NFAT1 in nuclear (N) and cytoplasmic (C) fractions of effector WT (gray) and YAPcKO (green) T cells. Cytoplasmic GAPDH and nuclear TATA-binding protein (TBT) served as loading and fractional controls. Data are representative of 3 independent experiments. (B) Schematic representation depicting the multiple domains and YAP mutants used in the study. (C) Co-immunoprecipitation analysis of proteins with anti-IQGAP1 in YAPcKO T cells transduced with FL YAP, YAPΔTBD, YAP 5SA, YAP 4SA, or GFP control. Immunoblotted proteins included IQGAP1 and NFAT1. Data representative of three independent experiments (dots at right). Quantification of NFAT1 pulled down was normalized to the ratio of NFAT1 to IQGAP1 pulled down in the control. (D) Co-immunoprecipitation analysis of proteins with anti-FLAG in YAPcKO T cells transduced with FL FLAG-YAP, FLAG-YAP 5SA, or GFP control. Immunoblotted proteins included FLAG-YAP and NFAT1. Data representative of two independent experiments (dots at right). Quantification of NFAT1 pulled down was normalized to the ratio of NFAT1 to FL FLAG-YAP. (E) YAPcKO T cells were transduced with human FL YAP, YAPΔTBD, YAP 5SA, YAP 4SA, or GFP control, sorted for GFP positive cells, and then stained for immunofluorescence. Representative images show NFAT1 localization for each condition. Scale bar, 2 μm. (F) Quantification of NFAT1 nuclear localization completed by ImageJ. Each data point represents a single cell, n > 200 per condition. Data representative of 3 independent experiments. Box shows mean and 95% CI. (G) Flow cytometric analysis of CD25 expression in WT and YAPcKO T cells treated with differential concentrations of cyclosporin A (CsA). Data from 3 independent experiments are shown. (H) Flow cytometric analysis of CD98 expression in WT and YAPcKO T cells treated with CsA as in (F). Data from 3 independent experiments are shown.

IQGAP1 is a cytoplasmic scaffold protein that interacts with over 100 partner molecules. In T cells, IQGAP1 sequesters NFAT1 in the cytoplasm and participates in regulating its transit to the nucleus ^33^. IQGAP1 was recently shown to interact with YAP and regulate its nuclear translocation as well ^34^. We hypothesized that in T cells YAP could regulate the ability of IQGAP1 to bind to phosphorylated NFAT1, and thus regulate NFAT1 nuclear translocation. We made mutants of YAP (**Fig. 5B**) and tested their interactions with IQGAP1 and their effects on NFAT translocation. We found that significantly more NFAT1 binds to IQGAP1 in YAPcKO T cells expressing FL YAP as compared to a control vector, suggesting that the presence of YAP augments the interaction between IQGAP1 and NFAT (**Fig. 5C**). In the canonical Hippo pathway that regulates YAP, the LATS1/2 kinases phosphorylate YAP to sequester it in the cytoplasm. We mutated four LATS1/2-targeted serine sites to alanine (YAP 4SA). Another modification that controls YAP activation is O-linked glycosylation, which is required for nuclear transport ^35^, of one of the 4SA serines. YAPcKO T cells expressing YAP 4SA had significantly more NFAT1 associated with IQGAP1 compared to those expressing the control vector, and similar levels of bound NFAT1 as T cells expressing FL YAP (**Fig. 5C**). This result shows that YAP influences the IQGAP-NFAT interaction, and this effect is not regulated by the canonical Hippo pathway or O-linked glycosylation in T cells. We further mutated serine 384 into alanine in addition to the mutations in YAP 4SA and termed this mutant YAP 5SA. This serine had been shown to be regulate degradation of YAP in other systems ^36^. In contrast to YAP 4SA, YAPcKO T cells expressing YAP 5SA failed to keep NFAT1 coupled to IQGAP1 (**Fig. 5C**). This result shows that serine 384 of YAP is essential to regulating NFAT activity through IQGAP1. The amino terminal, TEAD-binding domain (TBD) plays a critical role in the transcriptional regulation enacted by YAP in other systems, but the TBD is not necessarily required for all YAP-associated biology ^19^. To explore whether the TEAD-binding domain of YAP was required for IQGAP regulation of NFAT1, we expressed YAPΔTBD in YAPcKO T cells and found that there were similar amounts of phosphorylated NFAT1 bound to IQGAP1 as compared to T cells transduced with FL YAP or YAP 4SA (**Fig. 5C**). Thus, the YAP TBD is not required for regulating IQGAP1-NFAT1 interactions. Taken together, these results show that YAP regulates the interaction of IQGAP1 with NFAT1 in T cells, and that Ser384 of YAP plays a central role.

To assess whether YAP could inhibit NFAT1 nuclear localization we transduced the YAP constructs or control vector into YAPcKO T cells and then imaged for NFAT, YAP, and nuclear DNA. In YAPcKO cells expressing FL YAP, YAP 4SA, or YAPΔTBD, NFAT was excluded from the nucleus to a similar degree (**Fig. 5D-5E**). In contrast, we saw no restriction of NFAT localization to the nucleus with the YAP 5SA construct, which was similar to cells transduced with the control vector. Thus, YAP regulates sequestration of NFAT from the nucleus, and YAP Ser384 is important for this interaction. Despite the potential role of YAP Ser384 to regulate degradation in other systems, we saw no notable differences in YAP expression between YAP 4SA and YAP 5SA expressed in YAPcKO T cells (**Fig. 5D** and **Fig. S6B**). Thus, the trafficking of NFAT to the nucleus in T cells depends on Ser384 in YAP, but not the TBD, or the four serine phopshorylation sites in YAP 4SA. We assessed the biological impact of these YAP constructs and their regulation of NFAT on the suppression of proliferation of CD4+ T cells. YAPcKO T cells transduced with FL YAP or YAPΔTBD (i.e., those with intact Ser384) proliferated less than cells transduced with control vector or YAP 5SA (**Fig. S6C and D**). These results offer biological validation of the impact of YAP and its Ser384 in regulating NFAT1.

We questioned whether the role YAP played in suppressing T-cell responses was primarily mediated through NFAT. YAP in other systems regulates transcription through interaction with TEADs factors in the nucleus. In T cells, however, YAP-mediated suppression of responses was associated with cytoplasmic, not nuclear, localization (**Fig. S6B**). Furthermore, deletion of the TBD of YAP did not affect interaction with IQGAP1, translocation of NFAT to the nucleus, translocation of YAP to the nucleus, or proliferative responses of T cells. To identify any potential non-NFAT influence on YAP, we employed a strategy to modestly suppress NFAT while comparing activation of T cells from WT and YAPcKO mice. If YAP played an inhibitory role aside from on NFAT, we would expect to see WT T cells inhibited beyond YAPcKO T cells at all levels of NFAT suppression. We stimulated T cells using a modest antigen dose to allow for suppression, co-cultured with calcinuerin inhibitor cyclosporine-A (CsA). In the absence of CsA, YAPcKO T cells showed greater activation than WT controls. At low doses of CsA (10-25 nM), while activation was mostly still intact, YAPcKO T cells showed a diminishment of CD25 and CD98 expression to match the levels of WT effector T cells (**Fig. 5F-5G**). Above this dose, we observed that WT and YAPcKO T cells showed a lockstep decrease in effector responses, down to complete suppression of activation at high CsA. Any YAP-dependent, NFAT1-independent suppression of T-cell activation should not allow this effect. Thus, these data indicate that YAP suppresses T-cell effector responses through an NFAT-dependent mechanism.

### Sensing mechanical forces requires YAP

YAP is established as a key convergence point for mechanosignaling in many, and to test whether T cells sense mechanics through YAP, we returned to the system developed in **Fig. 1** and first examined whether substrate stiffness influenced NFAT nuclear localization. We activated SMARTA T cells with peptide-pulsed APCs atop soft (4 kPa) or stiff (40 kPa) scaffolds. We found significantly increased nuclear NFAT1 in T cells activated on the stiff surface as compared to soft, suggesting that mechanical forces supervise NFAT1 activity (**Fig. 6A**). In addition, CD25 and CD98, targets of NFAT, were upregulated in T cells activated on stiff surfaces compared to that of soft surfaces (**Fig. 6C**).

**Fig. 6.**
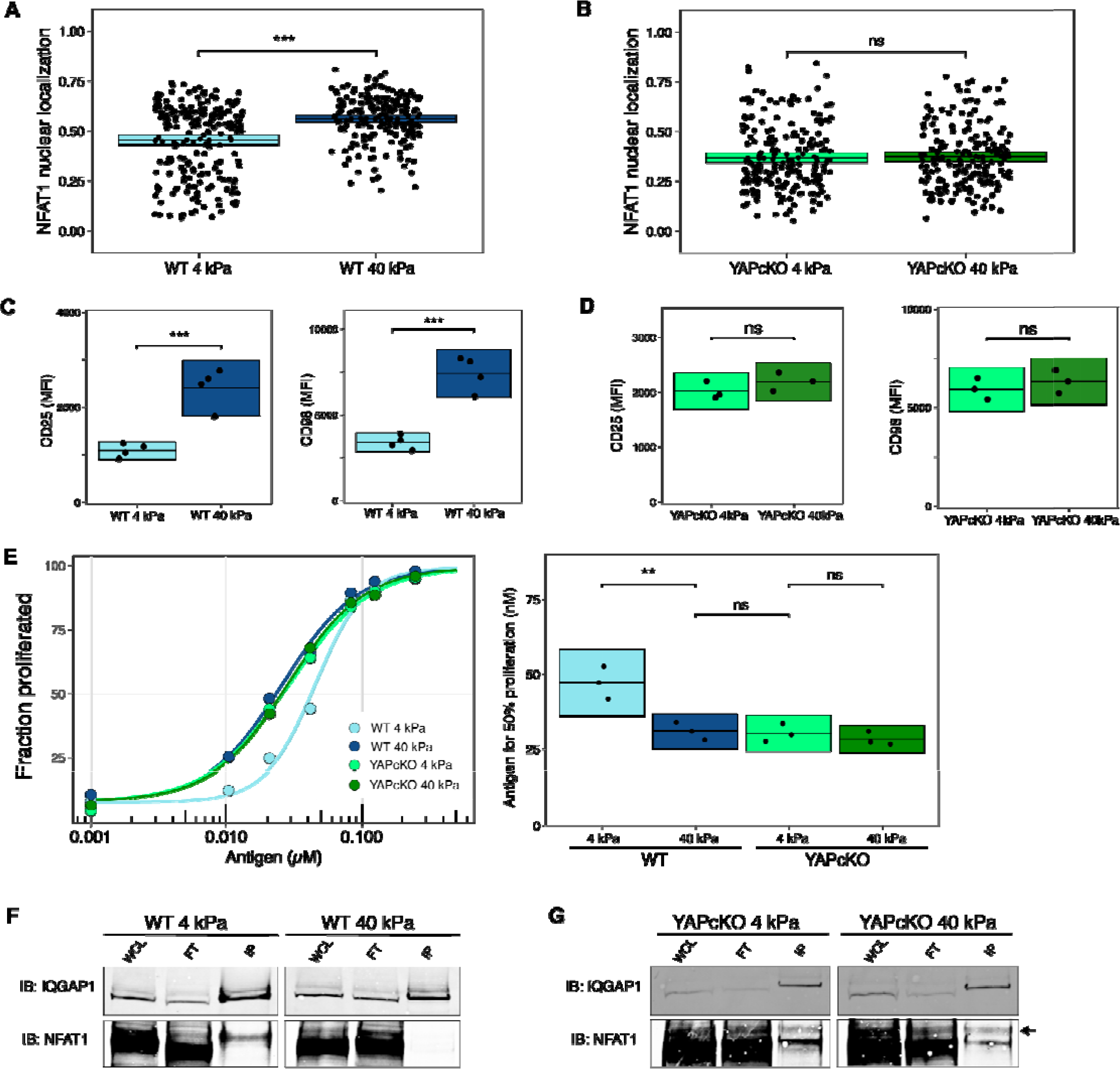
Substrate rigidity potentiates NFAT activity and cellular bioenergetics through YAP signaling. (A, B) SMARTA (A) WT or (B) YAPcKO T cells activated on soft (4 kPa) and stiff (40 kPa) alginate-RGD hydrogels were stained for NFAT1 for microscopy. Each dot represents a single T cell, n > 200 per condition, box shows mean and 95% CI. Data are representative of three independent experiments. (C, D) CD25 and CD98 expression of (C) WT or (D) YAPcKO T cells activated on soft and stiff alginate-RGD matrices. Box plot of CD25 expression for WT T cells was taken from data in **Figure 1D**. Each dot shows one independent experiment, box shows mean and 95% CI. (E) Dose response curve of GP 61-80 peptide with CFSE labeled SMARTA WT or YAPcKO T cells seeded onto soft or stiff 2D scaffolds with splenocytes. The amount of peptide required for 50% proliferation (ED50) from each dose response curve was calculated from 3 independent experiments and plotted. Box plot of ED50 for WT T cells was taken from data in **Figure 1C**. Data shown are from 3 independent experiments. Box shows mean and 95% CI. (F) Co-immunoprecipitation analysis of IQGAP1 in WT SMARTA T cells activated on soft and stiff substrates. Immunoblotted proteins included IQGAP1 and NFAT. Data representative of 2 independent experiments. (G) Co-immunoprecipitation analysis of IQGAP1 YAPcKO SMARTA T cells activated on soft and stiff substrates as in (F). Arrow indicates phospho-NFAT1 band. Data representative of 2 independent experiments.

These data led us to hypothesize that the mechanical microenvironment controlled NFAT activity in T cells through YAP. We compared NFAT1 nuclear/cytoplasmic expression in YAPcKO SMARTA T cells activated on soft and stiff surfaces. Importantly, the effect of elastic modulus on NFAT1 localization was entirely abolished in YAPcKO T cells (**Fig. 6B**). Additionally, YAPcKO T cells exhibited similar expression of NFAT1 targets CD25 and CD98 when exposed to stiff and soft microenvironments (**Fig. 6D**). These results indicate that NFAT activity and its downstream targets are dependent on YAP in T cells. YAPcKO T cells demonstrated similar proliferation kinetics to WT T cells on stiff substrates, further suggesting that mechanics-dependent T-cell proliferation is mediated by YAP (**Fig. 6E**). To test if substrate stiffness had a direct effect on the amount of YAP bound to IQGAP1 to control NFAT activity, we recovered T cells from soft and stiff scaffolds after activation and co-immunoprecipitated proteins complexed with IQGAP1. When we probed for the presence of NFAT1, we discovered that less NFAT1 was bound to IQGAP1 in T cells seeded on stiff hydrogels versus in T cells seeded on soft hydrogels (**Fig. 6F**). In contrast, NFAT1 was found complexed with IQGAP1 to a similar degree in YAPcKO T cells regardless of substrate stiffness (**Fig. 6G**). Thus, the mechanical environment influences NFAT nuclear localization, activation markers, proliferation, and IQGAP1-NFAT interaction in a YAP-dependent manner.

Activated T cells leaving an activated LN may find themselves circulating into LNs where autoantigens mimic a cognate stimulus. Molecular mimickry is indeed postulated to be a major driver of autoimmunity that becomes primed during infections and arises clinically shortly thereafter, for example, Type 1 diabetes. Our model would suggest that the mechanical softness of the unactivated LN should inhibit activated T cells bearing YAP and suppress autoimmunity. To test this hypothesis, we assessed whether self-reactive YAP-deficient CD4+ T cells could ignore the soft, inhibitory mechanical cues of naïve LN *in vivo*. We measured the ability of YAP to regulate Type 1 diabetes in NOD.SCID mice, which lack functional T and B cells. T cells bearing the islet-specific TCR BDC2.5 are diabetogenic when transferred into NOD.SCID mice, an outcome mediated by production of IFN-□ by activated T cells ^37^. We completed YAP knockdown by retrovirally transducing shRNA in BDC2.5 cells and verified that YAP protein was depleted (**Fig. S6E**). We adoptively transferred T cells transduced with YAP shRNA or control shRNA into NOD.SCID mice (**Fig. 6G**). We found that mice receiving T cells depleted of YAP developed diabetes faster than with those receiving control T cells (**Fig. 6I**). YAP-depleted BDC2.5 T cells developed into more IFN-□+ T cells on a per-cell basis than controls (**Fig. 6H**), proliferated to a greater extent in response to cognate peptide, and made more IL-2 (**Fig. S6F**). These results indicate that sensing the mechanical microenvironment in LN requires YAP to slow the progression of autoimmune disease. Taken all together, our data demonstrate the critical role of YAP as a mechanosensor that links tissue mechanics to regulation of the T-cell effector response through NFAT sequestration and metabolic control.

## DISCUSSION

Here, we identify a new physiological feedback mechanism whereby microenvironmental cues guide T-cell effector responses. These mechanical signals are separate from the mechanical forces exerted on the TCR, and thus T cells appear to be capable of independently sensing forces at both the molecular scale and tissues scale. The pathophysiological implication of these findings may extend from infections to autoimmunity to cancer, each of which shows considerable alterations in tissue mechanics. In this paper, we study autoimmunity and infection models, and show dramatic changes in the mechanics of the lymph nodes. Mechanical changes in LNs occur early due to migrating, activated dendritic cells that induce alterations in fibroblastic reticular cells ^38,39^, and later from proliferation of lymphocytes within the encapsulated, confined environment. We describe a YAP-mediated pathway by which a mechanically stiff microenvironment enhances T-cell activation and metabolic reprogramming as YAP enters the nucleus and allows for NFAT activation. As activated T cells emigrate from the LN and the pathogen is cleared, the stiffness of LNs starts to fall. In the mechanically soft microenvironment, YAP restrains T-cell activation and metabolic reprogramming as YAP enforces sequestration of NFAT in the cytoplasm by IQGAP1 (**Fig. S6E**). Thus, YAP controls the T cell’s ability to recognize the rising and falling micromechanical environment, thereby modulating the metabolic state accordingly. The sensing of mechanics is not strictly dichotomous, as tissue mechanics offer a continuum of rigidity. Our work thus identifies a new mechanical checkpoint by which the natural cycles of tissue mechanical changes directly fine-tune T-cell activation and cellular bioenergetics.

In studying the mechanical cues offered by natural tissues, our work quantified the mechanical state of LNs associated with acute viral infection. In reality, the microenvironment of activated LNs and tissues comprises not only changes in stiffness, but also inflammatory cytokines, dendritic cells and their costimulatory potential, fibroblastic reticular cells and their chemokine expression, and more. Lacking an experimental approach to influence only mechanics of tissues but not these other consequences of innate immune activation led us to choose a combination of *in vivo* and *in vitro* approaches in this paper.

A key finding in this paper is the mechanism by which YAP influences NFAT trafficking to the nucleus, which differs from the other ways that YAP enacts its roles ^40^. In resting T cells, IQGAP1 holds NFAT in a complex that includes many other partners ^33^. We show here that YAP enforces sequestration of NFAT by controlling IQGAP1’s ability to bind to NFAT. Importantly, the regulation of NFAT on mechanically defined substrates requires YAP. Our work offers potential follow up: IQGAP1 is known to regulate actin cytoskeleton dynamics in T cells ^41^ and many other cells ^42^. Future investigation into how environmental stiffness regulates IQGAP’s ability to control the actin cytoskeleton may shed more light into the mechanosignaling mechanisms of YAP in general.

YAP collects critical regulatory cues to identify environments bearing favorable metabolic substrates and mechanics. We show that YAP also plays a role in the highly motile cells of the immune system, providing an accelerator or brake on T-cell responses as they encounter infected and autoimmune tissue microenvironments. The results presented here offer clues of an orchestrated, molecular pathway allowing mechanical stiffness to modulate T-cell activation and metabolism. Our findings also newly reveal that the adaptive immune response is fine-tuned in accordance with physiological and pathophysiological changes in tissue mechanics.

## ACKNOWLEDGMENTS

We thank Fernando Camargo, Calvin Kuo, and Shane Crotty for mice; Jason Cyster for LCMV; the Bollyky lab for experimental help. We appreciate comments and suggestions from Julien Sage and Paul Bollyky.

### Funding

We acknowledge the Stanford Shared FACS Facility; the UCLA Jonsson Comprehensive Cancer Center (JCCC) Flow Cytometry Core Facility that is supported by National Institutes of Health awards P30 CA016042 and 5P30 AI028697; and the UCLA Metabolomics core. We acknowledge financial support from the NIH (R01 GM110482), the Stanford Child Health Research Institute, and the UCLA Children’s Discovery and Innovation Institute.

### Author Contributions

Conceptualization, Analysis, Writing: K.P.M. and M.J.B; Investigation, Visualization: K.P.M., F.J.M., and T.J.T; Funding Acquisition, Project administration, Supervision: M.J.B.

### Competing Interests

The authors declare no competing interests.

### Data and Materials Availability

All data are available in the main text or the supplementary materials. MTAs regulate the sharing of SMARTA and YAP^fl/fl^ mice.

